# Suppression of sost/sclerostin and dickkopf-1 promote intervertebral disc structure in mice

**DOI:** 10.1101/2021.07.01.449486

**Authors:** Tori Kroon, Paul Niziolek, Daniel Edwards, Erica L Clinkenbeard, Alexander Robling, Nilsson Holguin

**Affiliations:** Department of Biomedical Engineering, IUPUI, Indianapolis, IN, USA; Radiology & Imaging Sciences, IUPUI, Indianapolis, IN, USA; Indiana Center of Musculoskeletal Health, Indianapolis, IN, USA; Department for Anatomy and Cell Biology, IUPUI, Indianapolis, IN, USA; Department of Mechanical and Energy Engineering, IUPUI, Indianapolis, IN, USA

**Keywords:** Anabolic therapeutics, Chondrocyte and cartilage biology, Genetic animal models, Preclinical Studies, Wnt/β-catenin/LRPs

## Abstract

Intervertebral disc (IVD) degeneration is a leading cause of low back pain and characterized by accelerated extracellular matrix breakdown and IVD height loss but there is no approved pharmacological therapeutic. Deletion of Wnt signaling receptor Lrp5 induces IVD degeneration and suggests that Wnt signaling in the IVD may be responsive to inhibition of Wnt signaling inhibitors sost(gene)/sclerostin(protein) or dickkopf-1 (dkk1). Anti-sclerostin antibody (Scl-Ab) is an FDA-approved bone therapeutic that activates Wnt signaling. We (1) determined if pharmacological neutralization of sclerostin, dkk1 or their combination stimulate Wnt signaling and promote IVD structure and (2) determined the extent of the response of the IVD to global, persistent deletion of *sost*. Nine-week-old C57Bl/6J female mice (n=6-7/grp) were subcutaneously injected 2x/wk for 5.5 wk with scl-Ab (25 mg/kg), dkk1-Ab (25 mg/kg), 3:1 scl-Ab/dkk1-Ab (18.75:6.25 mg/kg) or vehicle (Veh). Separately, IVD of *sost* KO and WT (wildtype) mice (n=8, grp) were harvested at 16 weeks of age. First, compared to vehicle, scl-Ab, dkk1-Ab and 3:1 scl-Ab/dkk1-Ab similarly increased lumbar IVD height and β-catenin gene expression. Despite these similarities, scl-Ab decreased cellular stress-related heat shock protein gene expressions while neither dkk1-Ab nor scl-Ab/dkk1-Ab altered the same. Genetically and compared to WT, *sost* KO increased MRI-determined hydration and proteoglycan staining in the IVD. Notably, persistent deletion of *sost* was compensated by upregulation of *dkk1,* which consequently reduced the cell nuclear fraction for Wnt signaling transcription factor β-catenin in whole IVD. Lastly, RNA-sequencing pathway analysis confirmed the compensatory suppression of Wnt signaling and determined a reduction of cellular stress pathways. Together, suppression of sost/sclerostin or dkk1 each promote IVD structure by stimulating Wnt signaling, but sclerostin and dkk1 may differentially regulate cellular stress pathways. Ultimately, postmenopausal women prescribed scl-Ab injections to prevent vertebral fracture may also benefit from a restoration of IVD height and health.

**Graphical abstract:** 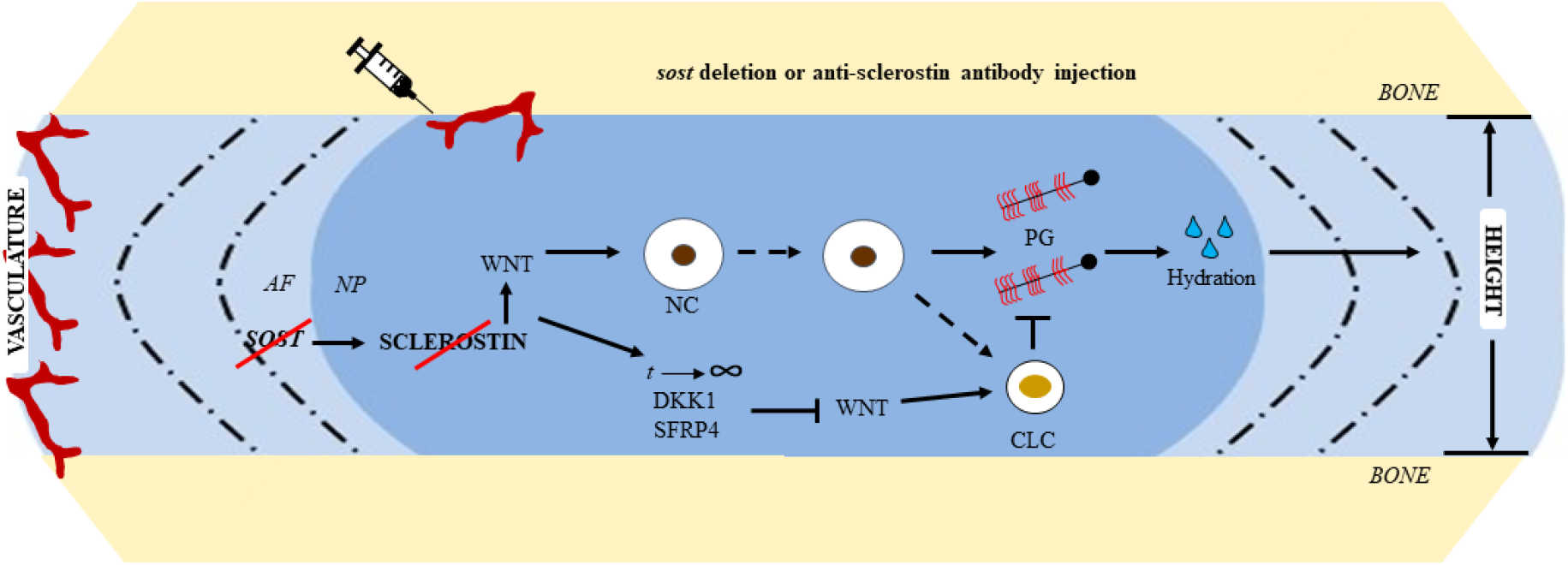

Suppression of Wnt signaling inhibitors by genetic or pharmacological approaches promotes intervertebral disc structure and hydration by Wnt signaling. However, persistent activation of Wnt signaling induces a compensatory reduction of Wnt signaling that shifts IVD cells toward a chondrocyte-like (CLC) phenotype. AF: annulus fibrosus, NC: notochordal cell, NP: nucleus pulposus, PG: proteoglycan

## Introduction

There are no FDA-approved pharmacological treatments for intervertebral disc (IVD) degeneration,^1,2^ a major contributing factor of low back pain.^3,4^ Osteoporosis may contribute to IVD degeneration^5^ and pharmacological treatments for bone maintenance may target the same anabolic pathways in the IVD. Anti-sclerostin-antibody treatment is an FDA-approved bone anabolic^6^ for postmenopausal women at high-risk of vertebral fracture.^7^ Sclerostin and dkk1 are inhibitors of the Wnt/β-catenin signaling pathway and global suppression of sclerostin by systemic injection of anti-sclerostin-antibody or genetic ablation of its precursor *SOST(human)/sost(mouse)* promotes bone formation and attenuates bone resorption.^8^ Individuals administered anti-sclerostin-antibody do not report an altered incidence of back pain than control subjects, suggesting it may be safe for the IVD.^6^ While osteocytes in bone are the major source of sclerostin^9,10^ and dkk1,^11^ IVD cells also express them,^12^ but the impact of sost or dkk1 regulation in the IVD has yet to be determined.

Sclerostin and dkk1 are inhibitors of the canonical wnt signaling pathway but differ in some notable ways. Both dkk1 and sclerostin interact with LRP5/6 to competitively prevent various Wnt ligands from binding to initiate the Wnt signaling pathway.^11^ Canonical Wnt signaling regulates cell fate and ECM anabolism in a range of musculoskeletal tissues. For instance, inactivation of Wnt signaling shifts differentiation of mesenchymal stem cells from osteoblastogenesis to chondrogenesis^13^ and activation in early chondrocytes triggers hyperchondrocyte maturation.^14^ Dkk1 primarily interacts with the third and fourth β-propellers of LRP5/6 but can bind to the first and second propellor,^15,16^ while sclerostin will bind with the first propellor.^17^ A pathway-related distinction between dkk1 and sclerostin is that dkk1 is a direct target of Wnt/β-catenin signaling pathway.^18^

In the spine, IVD development requires Wnt signaling,^19^ and loss of Wnt signaling by aging and IVD degeneration^20,21^ exacerbates ECM degradation.^20,22^ More specifically, notochordal cells of the centrally located nucleus pulposus require Wnt signaling to maintain their cell phenotype.^23^ Age- and injury-related reduction of Wnt signaling trigger the replacement of notochordal cells by chondrocyte-like cells.^20,24^ Contrarily, genetic stabilization of β-catenin in the nucleus pulposus, the hydration core of the IVD, increases notochordal cell expression and ECM anabolism^24^ and can prevent ECM degradation from a mechanical injury model of IVD degeneration.^22^ Lastly, in vivo deletion of LRP5 in IVD cells reduces Wnt signaling^20^ and suggests that the IVD may be sensitive to Wnt inhibitors that bind LRP5.

Therefore, we hypothesized that (1) neutralization of sclerostin and/or dkk1 and (2) deletion of Wnt signaling inhibitor *sost* would stimulate ECM anabolism in the IVD by increasing canonical Wnt signaling. Neutralization of sclerostin, dkk1 and in combination similarly increased Wnt signaling and IVD height. Next, using histology, MRI, qPCR and rna sequencing, global genetic deletion of *sost* increased the water content of the IVD, proteoglycan staining and decreased cellular stress mechanisms related to protein folding, but these changes were accompanied by gene and protein expression changes consistent with chondrogenesis by compensation of Wnt signaling. Overall, suppression of *sost*/sclerostin and/or dkk1 promote the structure of the IVD.

## Methods and Materials

### Mice

(1) 9-week-old C57Bl/6J female mice (n=6-8/group) were injected with either 25 mg/kg of sclerostin Antibody (SCL-AB), dkk1 antibody (DKK1-AB), a 3:1 ratio of the two antibodies (18.75 mg/kg of sclerostin Antibody and 6.25 mg/kg of dkk1 antibody), a 1:1 ratio (12.5 mg/kg of sclerostin and dkk1 antibody), a 1:3 ratio (6.25 mg/kg of sclerostin and 18.75 mg/kg of dkk1), or with buffer (VEH) (see Table 1) for 5.5 weeks, twice per week. (2) *Sost* KO mice and their wild-type (WT) littermates (n=8/group total) on a C57Bl/6 background have been previously described.^25^ Mice were housed in a 12-hour light/dark cycle, fed standard chow, and all experiments were performed with prior IACUC approval. Lumbar and caudal (tail) spinal sections were harvested. Mice were euthanized by hypoxia as a primary means and cervical dislocation as a secondary means. Spinal levels were further divvied-up for specific testing (Table 2).

**Table 1.** Scl-Ab/Dkk1-Ab dose by group

| Group | Scl-Ab (mg/kg) | Dkk1-Ab (mg/kg) | Total Ab (mg/kg) | n |
| --- | --- | --- | --- | --- |
| VEH | 0 | 0 | NA | 7 |
| SCL-AB | 25 | 0 | 25 | 6 |
| DKK1 | 0 | 25 | 25 | 7 |
| 3:1 | 18.75 | 6.25 | 25 | 7 |
| 1:1 | 12.5 | 12.5 | 25 | 7 |
| 1:3 | 6.25 | 18.75 | 25 | 7 |

**NA: Not applicable**

**Table 2.** Functional Spinal Unit for Each Outcome.

| Outcome | Tail | Lumbar |
| --- | --- | --- |
| MRI | CC6-7 | NA |
| Western Blot | CC6-8, CC10-12 | NA |
| mCT | CC7 | L6 |
| qPCR | CC8-10 | L3-5 |
| Histology | CC10-12 | L1-3 |

**NA: Not applicable**

### Histology and Immunohistochemistry

Injection group IVD as well as WT and *sost* IVD were run in a single batch. L1-3 and CC10-11 were fixed in 15 mL of 10% formalin on a rocker for 24 hours, submerged in 70% ethanol, embedded in paraffin, and sectioned (5 μm). Safranin-O/fast-green counterstain images were analyzed by 4 independent observers for an average IVD degeneration score.^20,26^ In short, the NP, AF and boundary between the two structures were scored based on structural properties and added for a total IVD score between 0-14, with increasing scores denoting greater IVD degeneration. Proteoglycan content in the NP was estimated as the amount of staining per area. Alcian blue staining was the counter stain for the IHC staining of sclerostin (BAF 1589, R&D Systems), osterix (#22552, Abcam), and collagen 2 (II-II6B3-c, DSHB). For both osterix and collagen 2 quantification in the NP, positive cells were counted and compared to the total number of cells that were stained brown for the protein and blue for the cell nuclei. For osterix quantification in the AF, the percent of the area-stained brown was measured and compared.

### Magnetic Resonance Imaging (MRI)

Motion segment CC6-7 was submerged and wrapped in 1x PBS-soaked gauze overnight until imaged. Imaging was completed on the Bruker BioSpin 9.4 T MRI, using a 400 nm slice thickness for 2D imaging. The motion segment was imaged in a sagittal orientation using a 0.052 × 0.052 mm voxel resolution in the x-y direction and a 0.4 nm voxel resolution in the z-direction taking 16 averages/slice. Two samples were stacked in a glass tube to remain upright, and two glass tubes were placed, separated by foam composite, inside of a 15mL tube to ensure samples would not move while being imaged. Images were analyzed for quantification by ImageJ (NIH). Area and intensity of the IVD were determined and multiplied to estimate the hydration content of the IVD.

### Micro-computed tomography

Motion segments L6-S1 and CC6-7 were harvested and submerged in 1x PBS prior to imaging. Specimens were imaged using the Bruker SkyScan 1272 Micro-CT at a resolution of 8 micrometers. Images of motion segment were contoured around the periosteal and the endosteal of the bone. For the trabecular analysis, the growth plate was used as a landmark and trabecular bone analysis consisted of the next 30 consecutive images (slices).^27^ For cortical analysis, the longitudinal center of the bone was identified and 15 images above and below were analyzed using the Bruker CTan64 MicroCT software.^16^ Parameters measured include BV/TV (bone volume/tissue volume), trabecular number (Tb.N), trabecular thickness (Tb.Th) for trabecular bone and cross-sectional thickness for cortical bone, using a lower threshold of 60 and upper threshold of 225 for analysis.

### QPCR

L3-5 and CC8-10 IVDs were harvested, frozen in liquid nitrogen, pulverized and suspended in TRIZOL (Ambion) until further processing.^16^ RNA isolation and purification steps were followed (RNeasy mini kit, Qaigen) and RNA concentration was quantified (Nanodrop). CDNA was synthesized (iScript, Biorad) from 400 ng of total RNA for the following Taqman probes (Life Technologies); *aggrecan* (Mm00565794_m1), *keratin8* (Mm04209403_g1), *dmp1* (Mm01208363_m1), *sost* (Mm00470479_m1), *adamts5* (Mm00478620_m1), *collagen1* (Mm00801666_g1), *collagen2* (Mm01309565_m1), *osterix* (Mm04209856_m1), *β-catenin* (Mm01350387_g1), *serpina1a* (Mm02748447_g1), *serpina1c*+ (Mm04207703_mH), *serpina1d* (Mm00842095_mH), *sostdc1* (Mm03024258_s1), *foxa2* (Mm01976556_s1), *axin2* (Mm00443610_m1), *sfrp4* (Mm00840104_m1), *gdf5* (Mm00433564_m), *hspa1b* (Mm03038954_s1), Cxcl9 (Mm00434946_m1), Il1b (Mm00434228_m1)Wnt16 (Mm00446420_m1), dkk1 (Mm00438422_m1), Wnt3a (Mm00437337_m1). Relative gene expression was normalized to IPO8 (Mm01255158_m1) for each group and then experimental values (*sost* KO) were normalized to the average of the WT value (2^−ΔΔCT^).

### Western Blotting

WT and *sost* KO IVDs between CC6-8 were isolated for whole cell lysate and cytoplasmic and nuclear separation Western Blots.^28^ IVDs were minced in ice-cold phosphate buffered saline (PBS; Fisher) containing 2% fetal bovine serum (FBS; Atlanta biologicals) and protease inhibitor PMSF (Sigma). Two tail IVDs from a single animal, per isolation method, were homogenized using a Tissue Tearor (BioSpec Products). Whole cell lysate was generated using diluted 1x cell lysis buffer (Cell Signaling) supplemented with PMSF. Fractionation of the nuclear protein was performed according to the Pierce cytoplasmic and nuclear extraction kit instructions (Sigma). Samples were run on an SDS-Page gel (BioRad) and transferred to PVDF membrane (BioRad). Blots were probed for anti-β-catenin (unphosphorylated; Cell Signaling) and subsequently an HRP-tagged anti-Rabbit secondary antibody (#7074S, Cell Signaling). Whole cell lysates were normalized to HRP-tagged Actin antibody (#A3854, Sigma Aldrich) and nuclear fractions to HRP-tagged Histone H3 (#12648, Cell Signaling). All blots were developed using Immobilon Luminata Forte (Sigma) and images collected digitally with the Amersham Imager 600 (GE Healthcare). Densitometry quantification was conducted on ImageJ to enumerate relative protein values between groups.

### Bioinformatics

To generate the top downregulated pathways in the *sost* KO, Gene Set Enrichment Analysis software, GSEA, was used. The input included gene name and raw counts for each sample. To generate the top up and downregulated pathways between the WT and *sost* KO group, WEB-based Gene SeT AnaLysis Toolkit, a freeware program, was used. Using GSEA method of interest and geneontology for the functional database and the input includes gene symbols and associated fold change for each gene. To generate plots such as PCA plot, volcano plot, and heat maps, R Studio was used.

### Statistics

A Dunnett’s test was used for the injection studies, with Veh for comparison, and a one sample t-test was used to test for outliers. A Student’s t-test was used between WT and *sost* KO IVD and between Veh and Scl-Ab. Box plots were used as the main graphical output. A box plot displays a 5-number summary of the data. The top and bottom lines of the box plot represent the first and third quartile while the median is the center line in the box. The top and bottom points represent the minimum and maximum of the data. A p-value or FDR-value (where applicable) < 0.05 was considered significant.

## Results

### Injection of neutralization for sclerostin, dkk1 and in 3:1 combination increased IVD height

Systemic administration of scl-Ab, dkk1-Ab, and the 3:1 combination of scl-Ab and dkk1-Ab increased lumbar IVD height by 22%, 26%, and 32%, respectively (Fig. 1A, B) but the 1:1 and 1:3 combination injections did not significantly increase IVD height. Neither IVD degeneration score nor proteoglycan staining intensity were altered by any injections. In tail IVD, no variety of injection impacted IVD height, proteoglycan staining intensity, or IVD degeneration score (Supp. 1).

**Figure 1.**
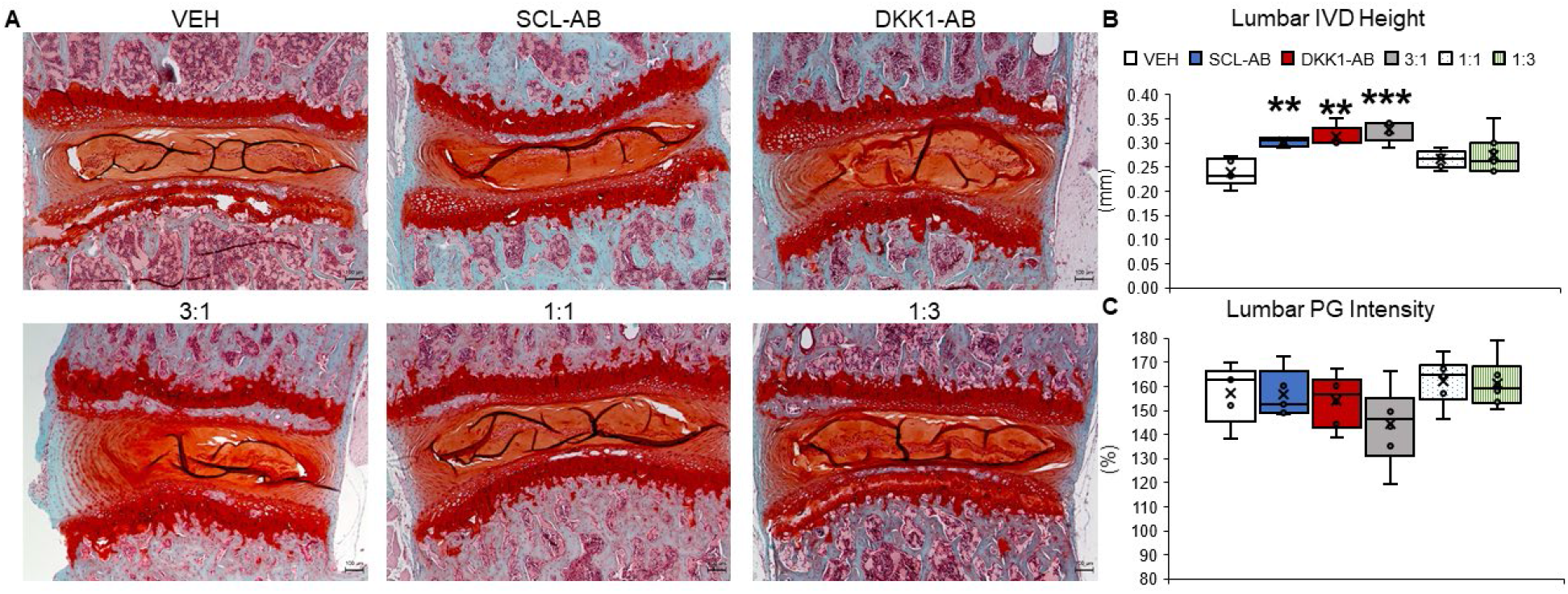
Qualitative and Quantitative Lumbar IVD Structural Measurements. (A) 5X magnification images of Safranin-O and Fast Green counter stained histological sections of the lumbar IVD for vehicle (VEH), 25 mg/kg sclerostin-antibody injection (SCL-AB), 25 mg/kg injection of dkk1-antibody (DKK1-AB) (top row, left to right), combination injection of 18.75 mg/kg sclerostin-antibody and 6.25 mg/kg dkk1-antibody (3:1), combination of 12.5 mg/kg each of sclerostin- and dkk1-antibody (1:1), and a combination injection of 6.25 mg/kg of sclerostin-antibody and 18.75 mg/kg of dkk1-anitbody (1:3) (bottom row, left to right). (B) Quantitative measurement of lumbar IVD height of all 6 groups. (C) Quantitative measurement of proteoglycan intensity staining in percentage of the lumbar IVD. Red staining indicating proteoglycan content. Scale bar is 100 μm. **p<0.01, ***p<0.001.

### Injection of sclerostin-neutralization-, dkk1-, and 3:1 combinatorial- antibody stimulated Wnt signaling

Quantitative PCR was determined in the injection groups that increased IVD height relative to vehicle: scl-Ab, dkk1-Ab and 3:1 scl-Ab:dkk1-Ab. Scl-Ab and dkk1-Ab upregulated transcription of *β-catenin* by 1.7-fold and 1.2-fold, respectively, while the 3:1 combination injection was trending toward upregulation (Fig 2A). *β-catenin* is a key transcriptional factor in the Wnt signaling pathway, where activation of this pathway allows for *β-catenin* to translocate to the nucleus to activate the expression of other target genes.^29^ Corroboratively, scl-Ab reduced *sost* gene expression by 85% and 3:1 combination reduced *sost* gene expression by 61%. Highly variable *dkk1* expression in the vehicle group obfuscated reduction by each injection group. Heat shock protein a1b (*hspa1b*) gene expression in the IVD was not different between groups but scl-Ab trended (p=0.069) to reduce *hspa1b*. *Hspa1b* and *serpinA1a* were similarly regulated by each antibody injection (R=0.83, p<0.05) but neither *hspa1b*, *serpina1a* nor *aggrecan* expression were changed by any injection compared to vehicle. Inflammation-related markers *cxcl9* and *il-1β* were not detected in any IVD.

**Figure 2.**
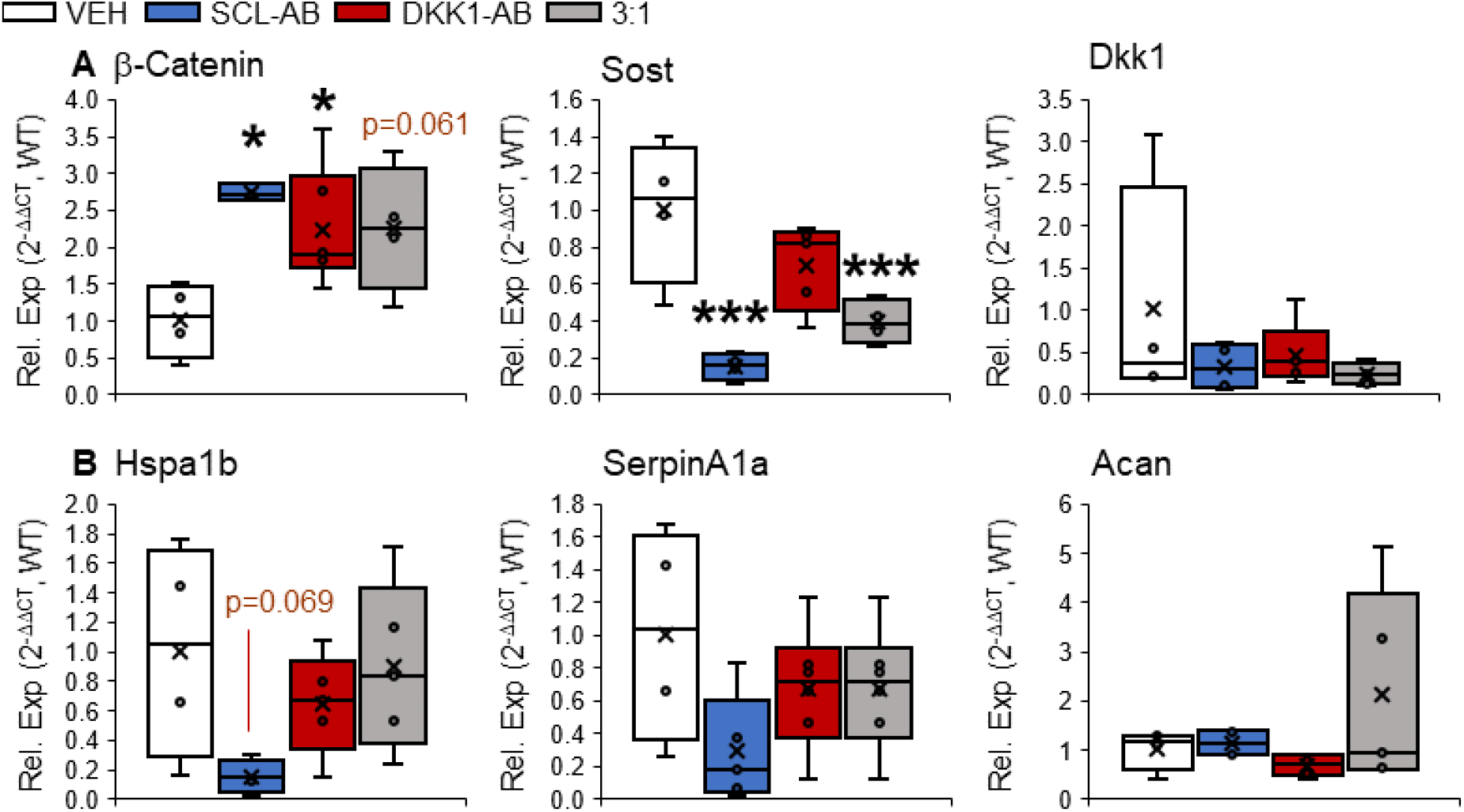
Injection of Scl-Ab and/or Dkk1-Ab increased Wnt signaling gene expression. (A) Gene expression related to Wnt signaling: β-catenin, Sost, and Dkk1. Gene expression of β-catenin in the 3:1 combination injection was trending toward significance (p=0.061) (B) Heat shock protein hspa1b, inflammatory marker serpina1a, and ECM marker aggrecan. Anti-sclerostin-antibody injection decreased gene expression of Hspa1b trending toward significance (p=0.069). *p<0.05, ***p<0.001.

### Sclerostin-antibody injection increased *β-catenin* protein and reduced chondrogenic gene expression in the IVD

Based on the low variability of upregulation of *β-catenin* by scl-Ab and trending downregulation of *hspa1b*, we further characterized ECM transcription and cell phenotype in the IVD by scl-Ab. Injection of scl-Ab decreased the gene expression of *col1a1* 56.7%, *col2a1* by 53.8%, and *osterix* by 34.7% (Fig. 3A). Further, sclerostin antibody increased the number of cells expressing β-catenin in the nucleus pulposus by 81% and reduced the number of nucleus pulposus cells that expressed *col2a1* by 46% (Fig. 2B, C). Similar to *hspa1b,* Scl-Ab decreased the gene expression of *hspa1a,* a gene associated with cellular stress^30^ by 95%.

**Figure 3.**
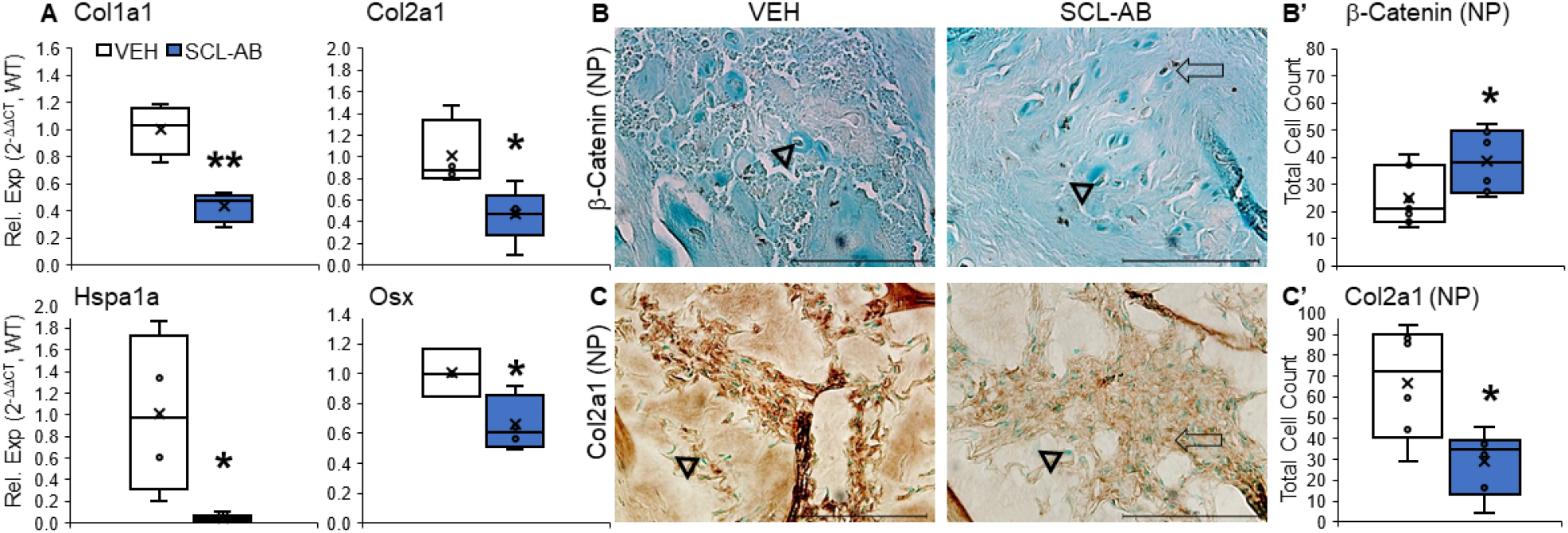
Anti-sclerostin antibody injection promoted β-catenin protein expression in the IVD and reduced chondrogenic gene expression. (A) Gene expression of anti-sclerostin antibody injection vs the vehicle group for *cola1a, col2a1, hspa1a,* and *osx.* (B) Images of β-catenin protein expression in the NP of the vehicle (VEH) and sclerostin antibody group (SCL-AB). (B’) Quantitative measurements of the fraction of β -catenin protein expressing cells to the total number of cells in the NP of both groups.

### Systemic deletion of *sost* reduced sclerostin gene and protein expression in the IVD

We analyzed sost KO mice to determine the impact of persistent reduction of sclerostin to the IVD. Quantitatively, *sost* gene expression was consistently detectable in the WT IVD but *sost* was not detected in any of the *sost* KO IVD (Fig. 4A). Qualitatively, sclerostin protein expression (brown, arrow) appeared in the NP cells of WT IVD and vertebral bone (Fig. 4B, Supp 2). By contrast, sclerostin staining in the *sost* KO was diffuse, minimally expressed in NP cell nuclei and relegated to the cell membrane. Osteocytes and osteoblasts are the predominate sclerostin-expressing cells.^29,31^ Here, WT tail vertebra expressed sclerostin staining in both osteocytes and osteoblasts while global deletion of *sost* blunted its expression in the vertebra (Supp. 2A) and, consequently, increased trabecular and cortical bone structure (Supp. 2B, C, Table 3).

**Figure 4.**
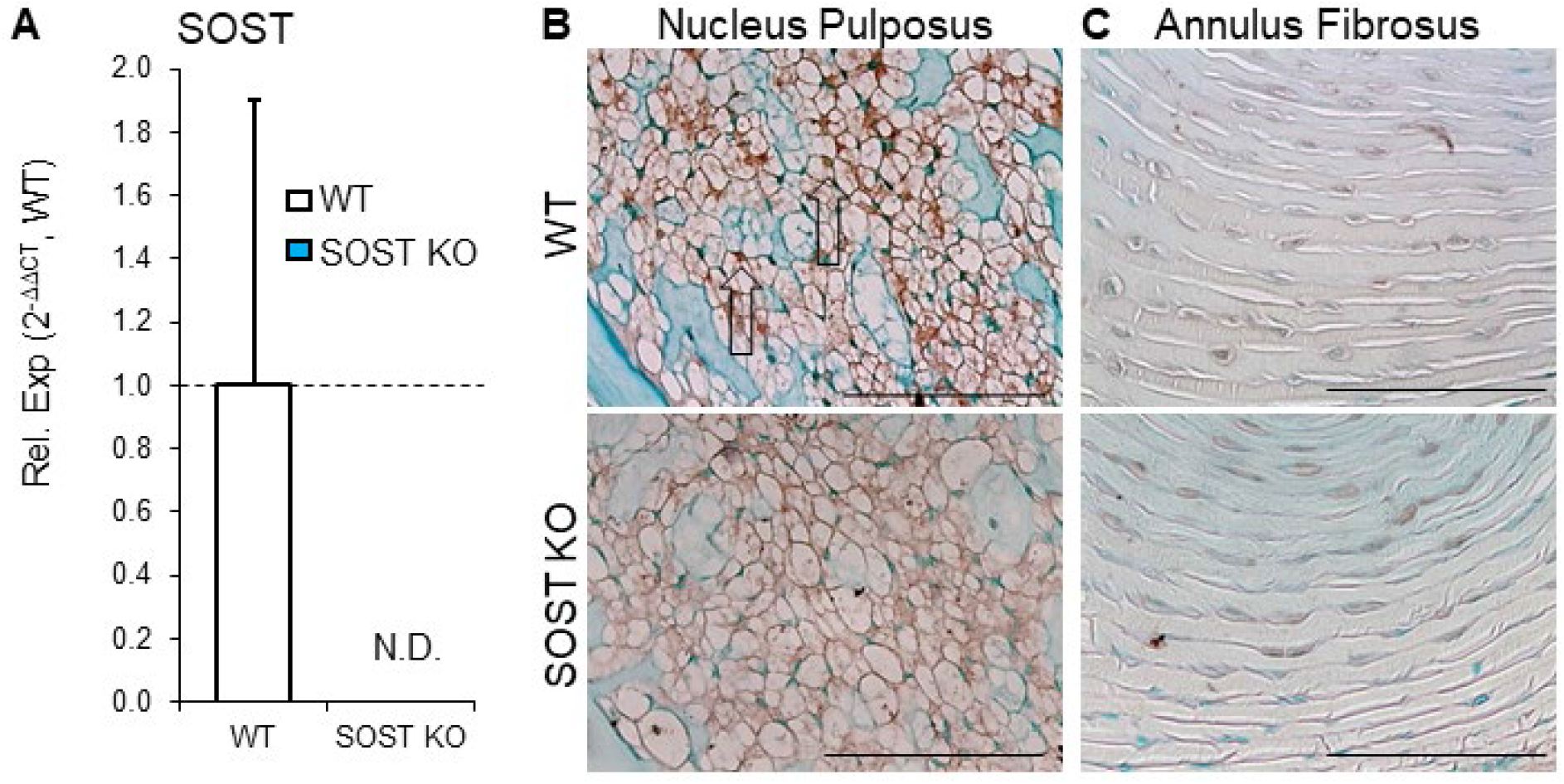
Sost deletion reduced gene and protein expression in the IVD. *Sost* gene expression from qPCR and qualitative images of sclerostin staining of the WT and *sost* KO IVD is effectively deleted from the *sost* KO group. (A) qPCR of *sost* with *sost* KO being normalized and compared to the WT. *Sost* gene expression was not able to be detected (N.D. – not detected). (B) 40X magnification images of the NP of the WT IVD (top) and *sost* KO IVD (bottom) using Alcian blue counterstain to show sclerostin protein expression in the cells of the IVD. The WT having more defined brown staining (black arrows) while the *sost* KO has less, more diffuse staining indicating less sclerostin expression in the *sost* KO. (C) 40X magnification image of the AF. Scale bar: 100um.

**Table 3.**
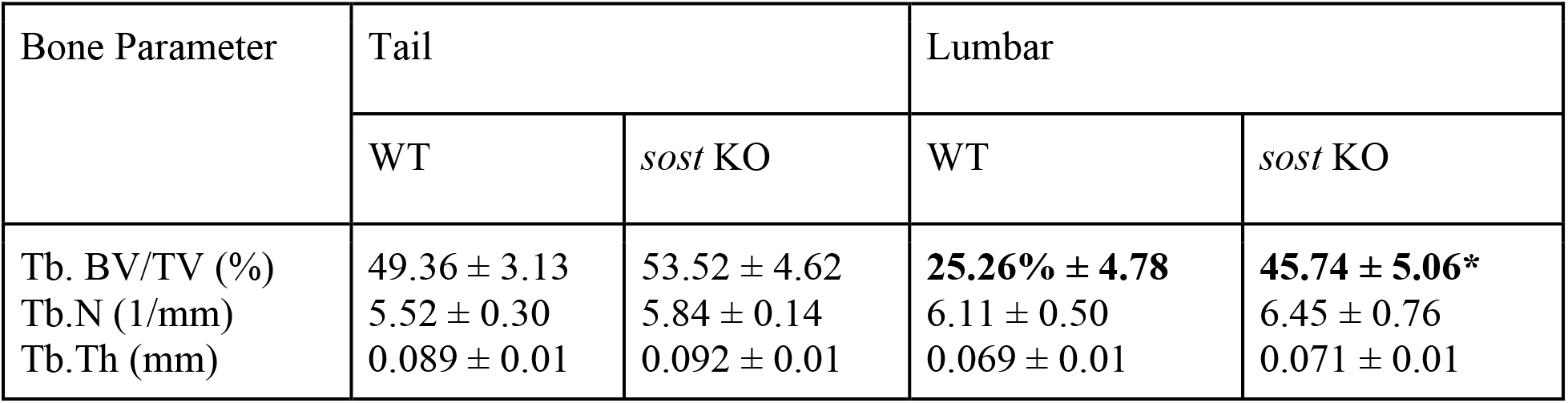
*sost* KO Vertebral Bone Structural Properties from uCT.

| Bone Parameter | Tail |  | Lumbar |  |
| --- | --- | --- | --- | --- |
|  | WT | *sost* KO | WT | *sost* KO |
| Tb. BV/TV (%) | 49.36 ± 3.13 | 53.52 ± 4.62 | **25.26% ± 4.78** | **45.74 ± 5.06\*** |
| Tb.N (1/mm) | 5.52 ± 0.30 | 5.84 ± 0.14 | 6.11 ± 0.50 | 6.45 ± 0.76 |
| Tb.Th (mm) | 0.089 ± 0.01 | 0.092 ± 0.01 | 0.069 ± 0.01 | 0.071 ± 0.01 |

### Deletion of *sost* in the IVD increased proteoglycan staining

The amount of proteoglycan, a crucial component to the IVD, can alter the overall structure of the IVD. WT IVD in both tail and lumbar have a singular band or mass of cells through the center of the NP. Compared to WT tail IVD, deletion of *sost* KO increased proteoglycan staining by 150% in the NP (Fig. 5A, B). However, the accumulation of proteoglycan staining, while potentially beneficial for hydration, led to slight disorganization of the band of cells in the NP, statistically insignificantly increasing the histological IVD degeneration score (Fig. 5A”). Tail IVD degeneration was associated with disorganization of the NP and lumbar IVD degeneration was associated with AF disorganization.

**Figure 5.**
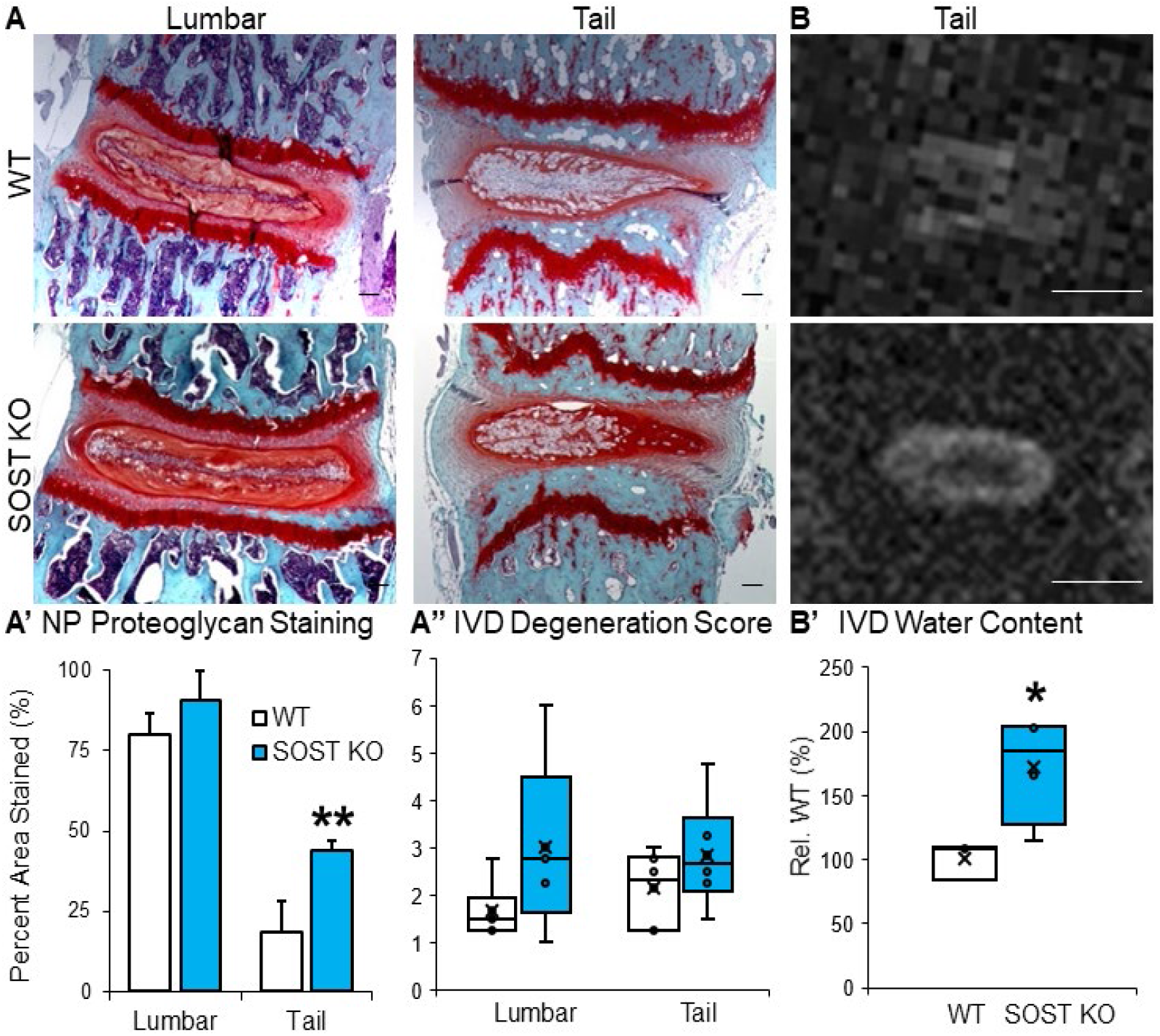
Deletion of *sost* promoted proteoglycan staining and IVD hydration. (A) 5X magnification images of Safranin-O and Fast Green counterstain of lumbar and tail IVD (left column, middle column) of the WT group (top row) and *sost* KO group (bottom row). Scale bar is 100 μm. (A’) Quantitative measurement of the intensity of proteoglycan staining in the IVD. Red staining indicating proteoglycan content. **p<0.01 (A”) Quantitative measurement of IVD degeneration score for lumbar and tai IVD of the WT and *sost* KO groups. (B) Using 9.4T MRI, tail motion segments were imaged highlighting the IVD of the WT (top) and *sost* KO (bottom). Scale bar is 1 mm. (B’) Relative water content quantitatively measured, using area multiplied by intensity, indicating increased hydration in the KO *p<0.05.

### Deletion of *sost* in the IVD increased IVD hydration

Proteoglycan is hydrophilic and deletion of sost increased proteoglycan staining in the NP. Therefore, we determined the hydration of the IVD by MRI, a standard non-invasive clinical imaging technique for determining the morphology and hydration of the IVD (Fig. 5).^32^ WT IVD had patched expression of water, lacking full concentric rings, but the ring of water that demarcated the cell band from the AF is clearer and brighter in the *sost* KO IVD (Fig. 5B). Relative to the WT IVD, *sost* KO increased the estimated water content, i.e., hydration, by 72% but there was no statistical difference in the total area of the IVD or intensity of the water between groups (Fig. 5B, B’, Table 4). These data show that deletion of *sost* may have imbibed the IVD.

**Table 4.** Area, Intensity, and Water Content Determined by MRI in WT and *sost* KO IVD

| Genotype | Area (mm<sup>2</sup>) | Intensity | Water Content |
| --- | --- | --- | --- |
| WT 1 | 0.32 | 956 | 301.12 |
| WT 2 | 0.32 | 1235 | 395.20 |
| WT 3 | 0.32 | 1004 | 391.56 |
| **WT AVG** | **0.34 ± 0.04** | **1065 ± 149.2** | **362.63 ± 52.29** |
| SOST KO 1 | 0.65 | 1145 | 739.67 |
| SOST KO 2 | 0.64 | 1155 | 733.43 |
| SOST KO 3 | 0.34 | 1225 | 416.50 |
| SOST KO 4 | 0.4 | 1501 | 600.40 |
| **SOST KO AVG** | **0.51 ± 0.16** | **1256.5 ± 166.8** | **622.50 ± 151.61\*** |

### *Sost* deletion induced compensation of Wnt signaling by upregulation of Wnt inhibitors

Canonical Wnt signaling requires translocation of β-catenin to the cell nucleus. Therefore, we determined the nuclear fraction of β-catenin. *Sost* KO IVD had less active β-catenin protein in the cell nuclei than WT IVD (Fig. 6A), suggesting *less* Wnt signaling. Dkk1 is a target of Wnt/β-catenin signaling so we determined the gene expression of Wnt inhibitors. Sost KO IVD expressed greater *dkk1* by 10-fold and *sfrp4* by 4-fold (Fig. 6B). Lack of gene expression difference of transcription factor *β-catenin* between WT and sost KO corroborated Wnt signaling compensation.

**Figure 6.**
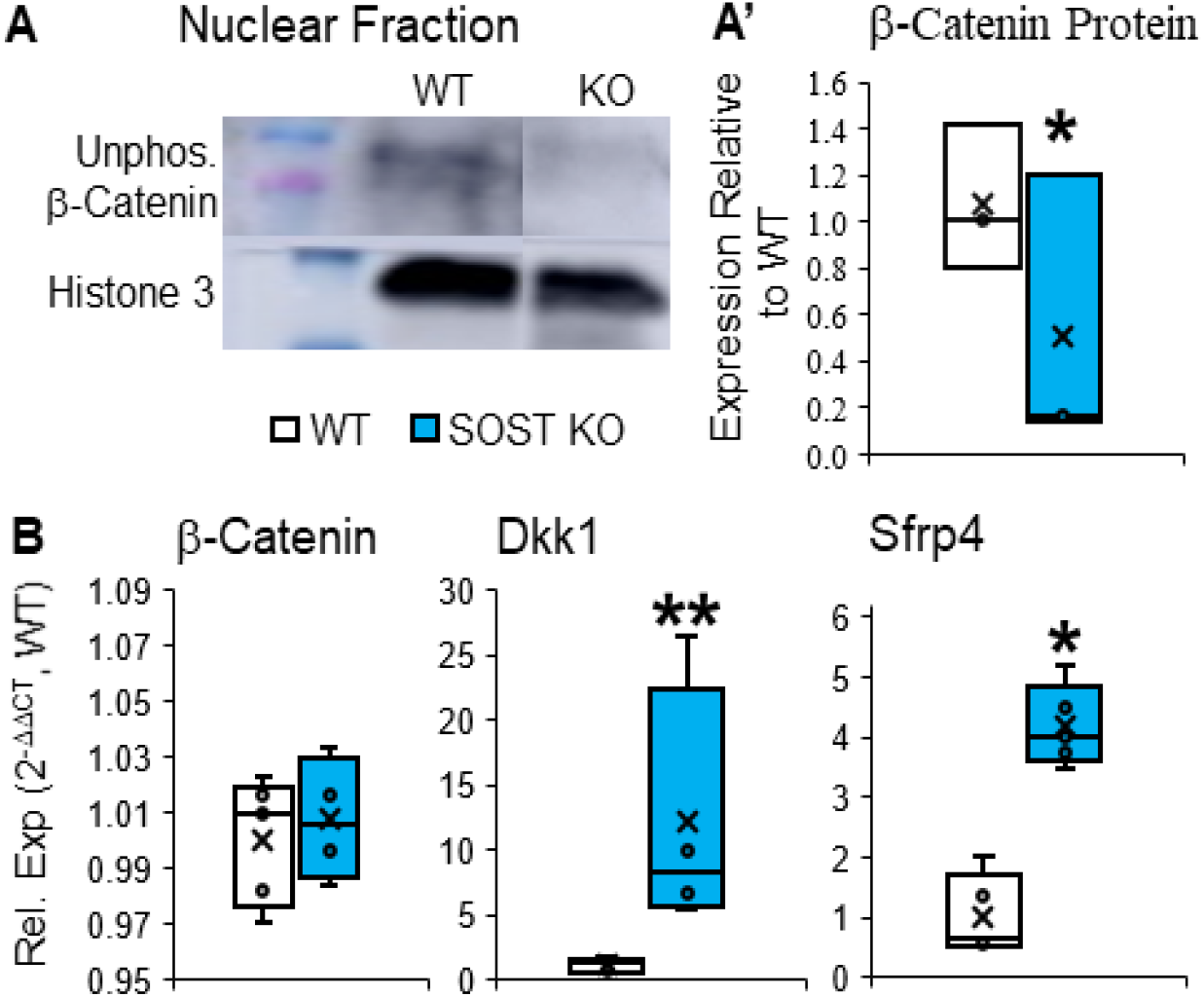
Deletion of sost induced compensation of Wnt signaling by upregulation of wnt inhibitors. (A) Nuclear separation western blot shows decreased amount of active β-catenin in the cell nuclei of the *sost* KO, using Histone 3 as control. (A’) Quantitative measurement of β-catenin protein expression from western blot. *p<0.05 Unphosphorylated (active) β-catenin at 92 kDa, Histone 3 at 17 kDa, β-actin at 92 kDa. (B) Genetic regulators of the Wnt signaling pathway. While β-catenin is unchanged, *dkk1* and *sfrp4*, both inhibitors of the pathway, are both increased in response to the *sost* KO. *p<0.05, **p<0.01.

### Compensation of Wnt signaling from *sost* deletion triggered extracellular matrix degradation and expression chondrogenic expression in the IVD

*Sost* KO regulated gene and protein expression of extracellular matrix metabolism towards anti-anabolism and catabolism and IVD cell phenotypes toward chondrogenesis. Specifically, deletion of *sost* downregulated gene expression of *aggrecan* (Fig. 7A).Next, common markers of notochordal cells, early chondrocyte-like cells (CLCs) and mature CLCs were determined, respectively. *Sost* KO IVDs expressed less transcription of notochordal marker *foxA2*^30^ by 77%, less transcription of early CLC marker *col2a1* by 1.6-fold, and fewer cells in the NP expressed *col2a1* (Fig. 7B, B’). By contrast, accrual of CLCs can be marked by increased chondrogenic marker^33^ and osteoblast transcription factor Sp7 (Osx). Compared to WT IVD, *sost* KO IVD expressed more osterix gene and protein expression (Fig. 7B, C). Sost deletion similarly altered gene expression in lumbar IVD with upregulation of ECM degradation, downregulation of notochordal marker *gdf5*^34^, downregulation of early CLC markers (*ColX*^35^) and upregulation of mature chondrogenic marker *bglap* (Supp 3). Gene expression that was not statistically different between WT and *sost* KO IVD included *serpinA1a-c* (*serpinA1c+*, p=0.06), *serpinA1d*, *sostdc1*, *keratin8*, *wnt16*, *axin2*, *serpinA1a*, *cxcl9*, *il-1β* and *wnt3a* (Table 3).

**Figure 7.**
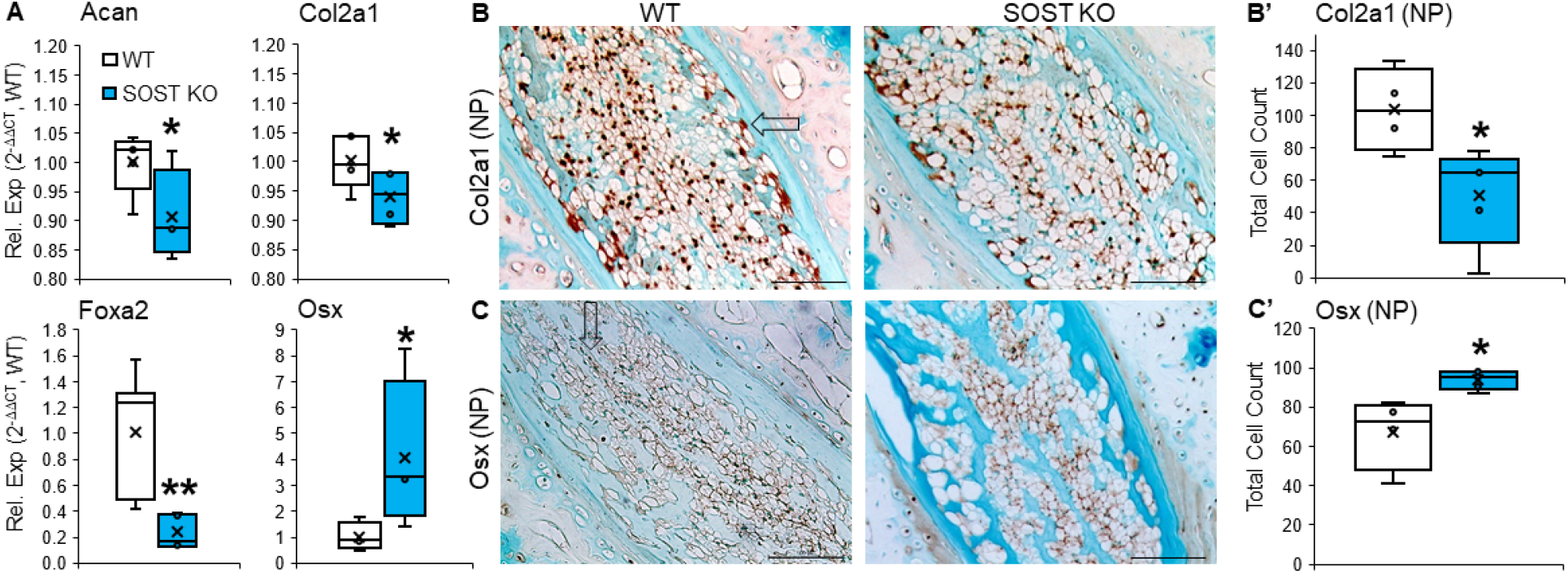
Global deletion of *sost* induced chondrogenic gene expression in the IVD. (A) Genes related to early chondrocyte-like cells (CLCs) *aggrecan* and *col2a1* (top row) are both decreased by the global *sost* deletion. *Foxa2,* a notochordal marker, and *osx,* a mature chondrocyte marker, are differentially expressed. *Foxa2* is decreased while *osx* is increased indicating a different expression profile in the *sost* KO IVD. (B) Collagen 2 staining of the NP of the WT and *sost* KO. (B’) Quantitative measurement of the fraction of the number of positively (brown) stained (black arrow) collagen 2 expressing cells. (C) Osterix staining of the NP of the WT and *sost* KO. (D) Quantitative measurement of the fraction of the number of positively (brown) stained Osterix expressing. *p<0.05, **p<0.01.

### Whole transcriptomic analysis of sost deletion in IVD show downregulated pathways related to protein folding and upregulated pathways related to immune response

The PCA plot (Fig. 8A) visualizes that the *sost* KO IVDs are not different, in an overall sense, to the WT IVDs. Nevertheless, the top 20 downregulated pathways were related to 35% cellular response to external stimuli (e.g., protein folding, FDR<0.05), 15% each metabolism and extracellular matrix organization, 10% gene expression, and 5% to developmental biology, signaling transduction, metabolism of proteins, transportation of small molecules, and other pathways (Fig. 8B). Whereas, the top 20 upregulated pathways were related to 85% immune (FDR<0.05), and 5% each cell cycle (FDR<0.05), signaling transduction (FDR<0.05), and hemostasis (FDR<0.05). Specifically, the heat map for the top downregulated pathway was ‘protein folding’ for cellular response, which included the highest number of heat shock proteins downregulated, and for the top upregulated pathway was the immune response (Fig. 8C). More specifically, the volcano plot highlights in red the genes that were most significantly differentially regulated (Fig. 8D). A list of the top 20 differentially downregulated and top 20 upregulated were included (Table 6, 7). Lastly, GSEA analysis for the *sost* KO IVDs corroborated that Wnt signaling was reduced based on the normalized enrichment score (−1.53, FDR<0.01, Table 8).

**Figure 8.**
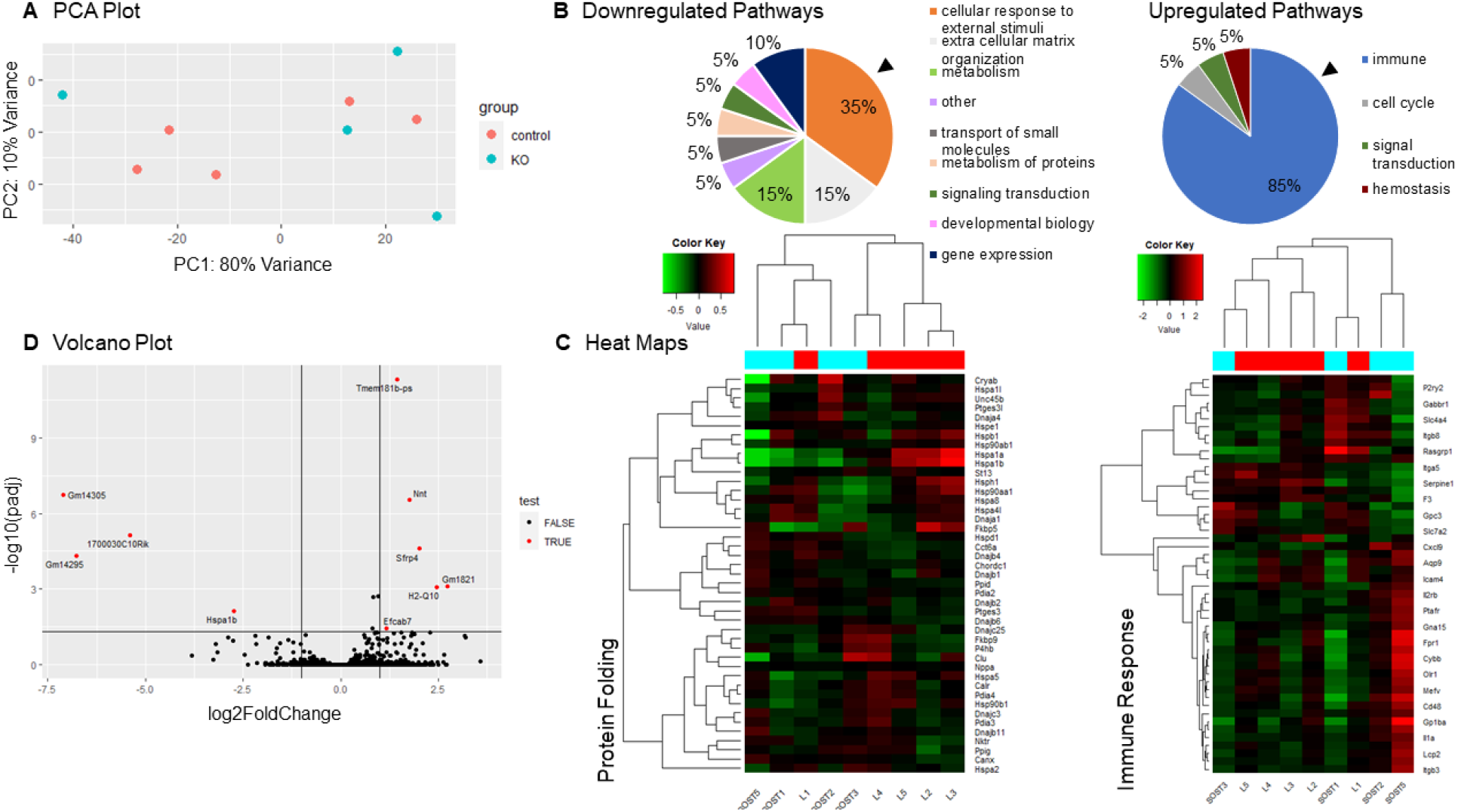
RNA sequencing of sost KO and WT IVD. (A) PCA plot of *sost* KO and WT IVDs demonstrating an overlap between the groups. (B) Top 20 downregulated and top 20 upregulated pathways following *sost* deletion. (C) Corresponding heat maps for protein folding associated with the top downregulated pathway (arrowhead in B) and immune response associated with the top upregulated pathway (arrowhead in B). (D) Volcano plot of all differentially regulated genes, with labels of most significantly regulated genes (FDR<0.05).

**Table 5.** Not Significantly Regulated Genes by Deletion of *sost* or Expressed Genes in the IVD.

| Gene | WT | *sost* KO | p-value |
| --- | --- | --- | --- |
| SerpinA1C+ | $1 \pm 0.18$ | $0.72 \pm 0.09$ | 0.06 |
| SerpinA1D | $1 \pm 0.56$ | $0.52 \pm 0.19$ | 0.11 |
| Sostdc1 | $1 \pm 0.35$ | $1.86 \pm 0.71$ | 0.11 |
| Keratin8 | $1 \pm 0.83$ | $0.50 \pm 0.50$ | 0.26 |
| Wnt 16 | $1 \pm 0.69$ | $1.79 \pm 1.46$ | 0.36 |
| Axin 2 | $1 \pm 0.53$ | $1.38 \pm 0.75$ | 0.38 |
| SerpinA1A | $1 \pm 0.37$ | $0.52 \pm 0.19$ | 0.80 |
| Cxcl9 | Undetermined | Undetermined | N/A |
| Il1b | Undetermined | Undetermined | N/A |
| Wnt 3a | Undetermined | Undetermined | N/A |

**Table 6.**
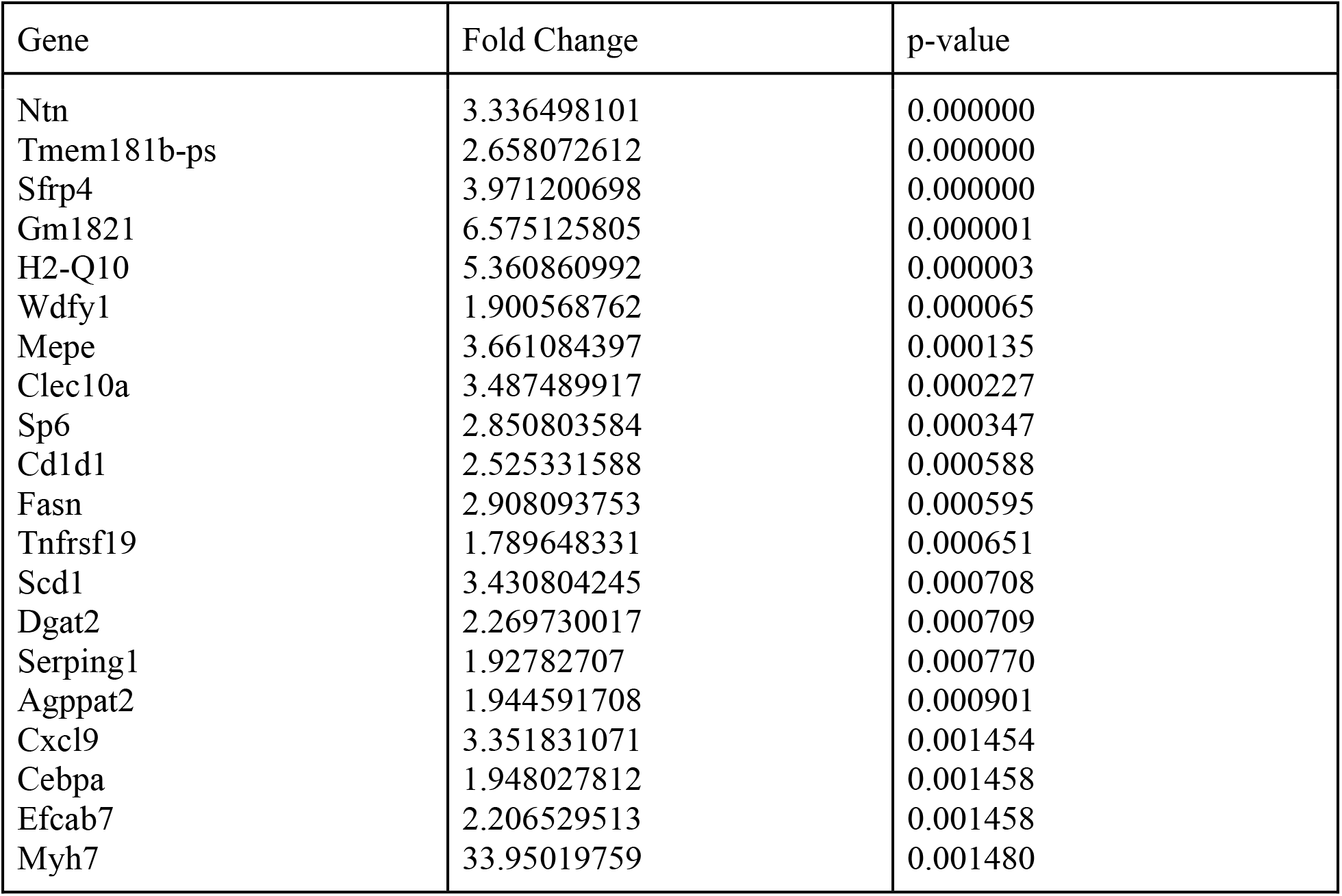
Top 20 Up-regulated Genes in *sost* KO Group.

| Gene | Fold Change | p-value |
| --- | --- | --- |
| Ntn | 3.336498101 | 0.000000 |
| Tmem181b-ps | 2.658072612 | 0.000000 |
| Sfrp4 | 3.971200698 | 0.000000 |
| Gm1821 | 6.575125805 | 0.000001 |
| H2-Q10 | 5.360860992 | 0.000003 |
| Wdfy1 | 1.900568762 | 0.000065 |
| Mepe | 3.661084397 | 0.000135 |
| Clec10a | 3.487489917 | 0.000227 |
| Sp6 | 2.850803584 | 0.000347 |
| Cd1d1 | 2.525331588 | 0.000588 |
| Fasn | 2.908093753 | 0.000595 |
| Tnfrsf19 | 1.789648331 | 0.000651 |
| Scd1 | 3.430804245 | 0.000708 |
| Dgat2 | 2.269730017 | 0.000709 |
| Serping1 | 1.92782707 | 0.000770 |
| Agppat2 | 1.944591708 | 0.000901 |
| Cxcl9 | 3.351831071 | 0.001454 |
| Cebpa | 1.948027812 | 0.001458 |
| Efcab7 | 2.206529513 | 0.001458 |
| Myh7 | 33.95019759 | 0.001480 |

**Table 7.**
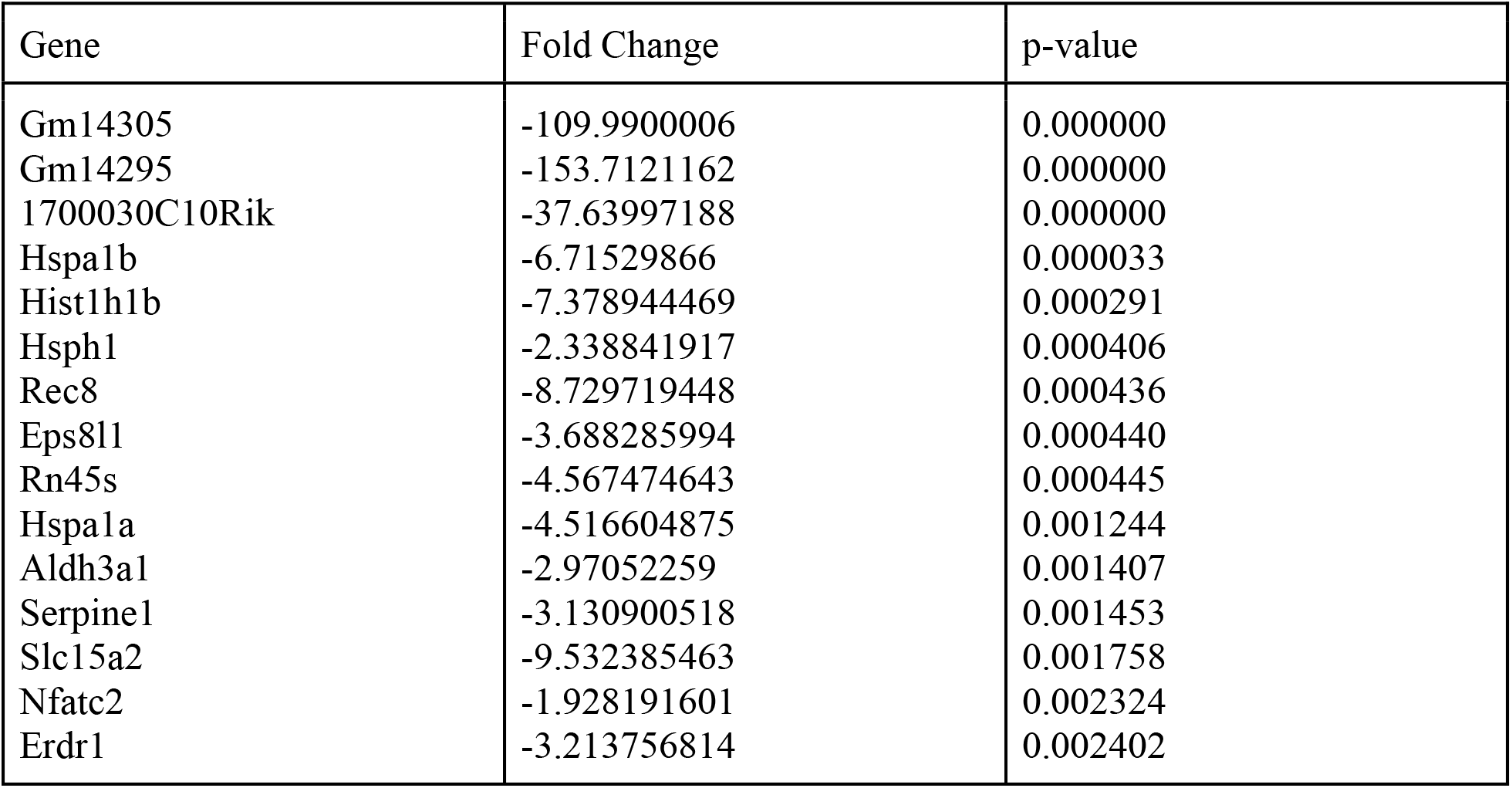
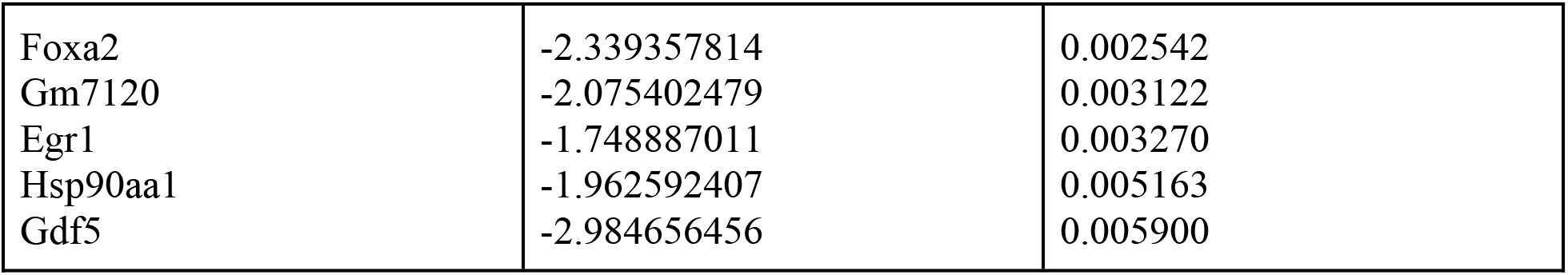
Top 20 Down-regulated Genes in *sost* KO Group.

**Table 8.**
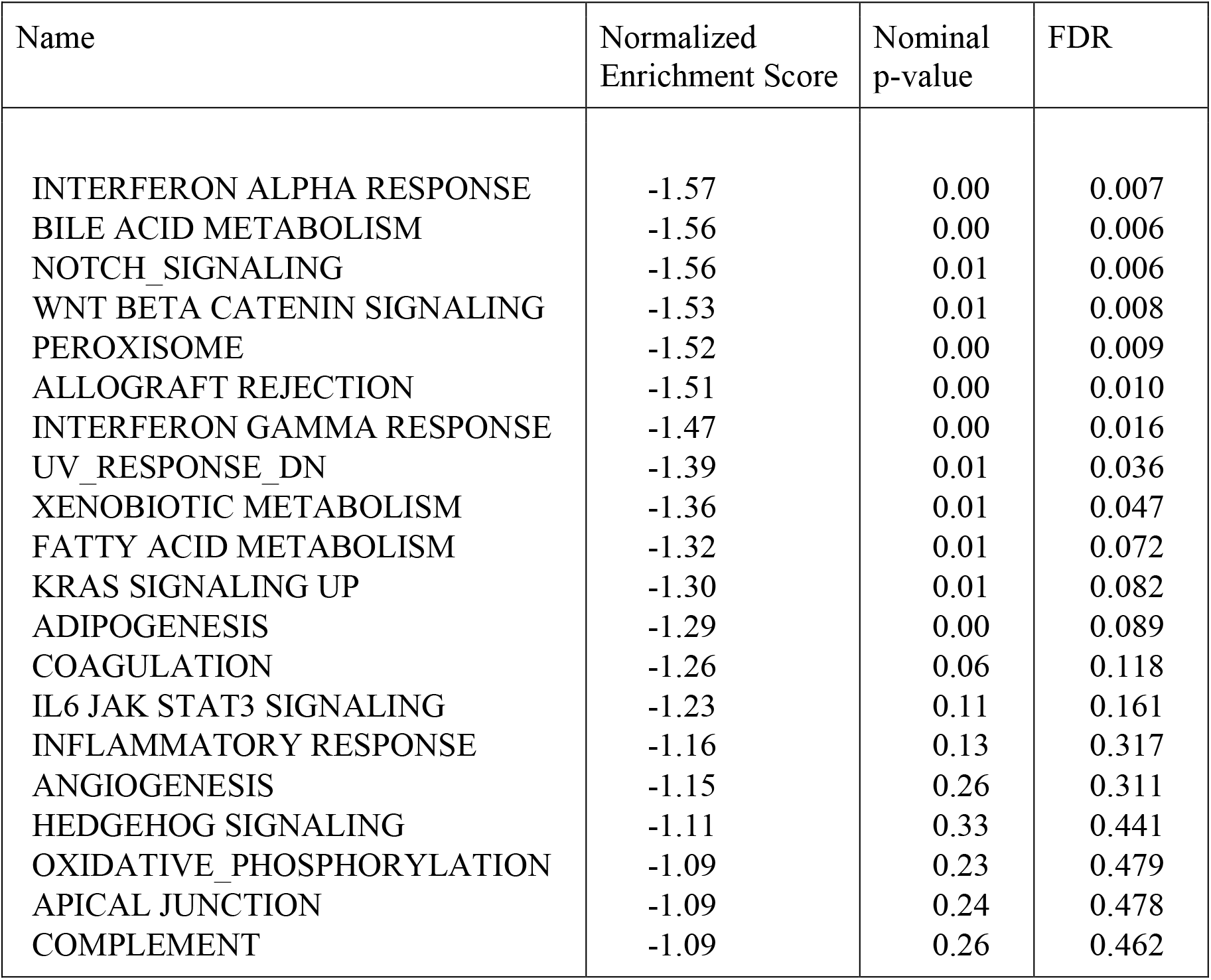
Top 20 downregulated pathways in *sost* KO group

| Name | Normalized Enrichment Score | Nominal p-value | FDR |
| --- | --- | --- | --- |
| INTERFERON ALPHA RESPONSE | -1.57 | 0.00 | 0.007 |
| BILE ACID METABOLISM | -1.56 | 0.00 | 0.006 |
| NOTCH_SIGNALING | -1.56 | 0.01 | 0.006 |
| WNT BETA CATENIN SIGNALING | -1.53 | 0.01 | 0.008 |
| PEROXISOME | -1.52 | 0.00 | 0.009 |
| ALLOGRAFT REJECTION | -1.51 | 0.00 | 0.010 |
| INTERFERON GAMMA RESPONSE | -1.47 | 0.00 | 0.016 |
| UV_RESPONSE_DN | -1.39 | 0.01 | 0.036 |
| XENOBIOTIC METABOLISM | -1.36 | 0.01 | 0.047 |
| FATTY ACID METABOLISM | -1.32 | 0.01 | 0.072 |
| KRAS SIGNALING UP | -1.30 | 0.01 | 0.082 |
| ADIPOGENESIS | -1.29 | 0.00 | 0.089 |
| COAGULATION | -1.26 | 0.06 | 0.118 |
| IL6 JAK STAT3 SIGNALING | -1.23 | 0.11 | 0.161 |
| INFLAMMATORY RESPONSE | -1.16 | 0.13 | 0.317 |
| ANGIOGENESIS | -1.15 | 0.26 | 0.311 |
| HEDGEHOG SIGNALING | -1.11 | 0.33 | 0.441 |
| OXIDATIVE_PHOSPHORYLATION | -1.09 | 0.23 | 0.479 |
| APICAL JUNCTION | -1.09 | 0.24 | 0.478 |
| COMPLEMENT | -1.09 | 0.26 | 0.462 |

## Discussion

FDA-approved anti-sclerostin antibody (scl-Ab) reinvigorated the structure of the IVD. We hypothesized that increasing Wnt signaling by suppression of Wnt inhibitors by pharmacological or genetic approaches would promote the structure of the IVD. Sclerostin and dkk1 are known inhibitors of the canonical Wnt signaling pathway and work in a similar fashion.^36^ Scl-Ab, dkk1-Ab, and the 3:1 combinatorial injection of sclerostin and dkk1 increased lumbar IVD height via strong upregulation of Wnt/β-catenin signaling but injection of a 1:1 ratio of sclerostin to dkk1 antibody or a 1:3 ratio were not as beneficial. Prolonged suppression of sclerostin by global deletion of *sost* increased water content and proteoglycan staining but induced a significant normalization of Wnt signaling. Overall, these data show that systemic administration of scl-Ab promotes major features lost with IVD degeneration.

### Increasing Wnt signaling by neutralization of sclerostin or dkk1 impose similar beneficial structural changes to the IVD

Osteocytes and osteoblasts are the dominant sclerostin-expressing cells in the body and inhibition of sclerostin or dkk1 induce bone formation.^37, 38^ IVD cells also express *sost*/sclerostin and dkk1^11^ but their role in the IVD is unclear. The Wnt/β-catenin signaling pathway plays a necessary role in IVD development but excessive activation of this pathway can disorganize IVD structure.^19^ Nucleus pulposus-specific upregulation and downregulation of Wnt signaling transcription factor β-catenin is anabolic and catabolic to the ECM of the IVD, respectively.^22^ Here, injections of scl-Ab or dkk1-Ab increased levels of β-catenin, lumbar IVD height and proteoglycan staining in the nucleus pulposus. IVD degeneration is characterized by IVD height loss from proteoglycan breakdown and dehydration,^40,41^ which is necessary to withstand spinal mechanical forces and function properly. Deletion of *sost* similarly increased the proteoglycan staining in the nucleus pulposus and increased water content, as measured by MRI.

### Compensation by sclerostin or dkk1 may serve to normalize Wnt signaling

Our data and others have noted that sclerostin and dkk1 share a mutual compensatory regulation in the Wnt/β-catenin signaling pathway. Previous studies have shown that dkk1 inhibition has strong anabolic effects on bone when sclerostin is disabled.^11^ We found that global sost deletion induced a strong upregulation of Wnt signaling inhibitors that normalized β-catenin protein in IVD cell nuclei and pathway analysis of Wnt signaling.

### Cell Phenotype

This suppression of Wnt signaling despite sost-deficiency induced an IVD cell phenotype switch to chondrogenic-expression that was, by contrast, suppressed in the injection studies.

In progress. . .

Despite imbibition of tail IVD and pro-anabolic ECM expression in lumbar IVD, canonical Wnt signaling was normalized in tail IVD and may have influenced the differences in structure and cellular phenotype between lumbar and tail IVD. Gene expression of Wnt signaling transcription factor β-catenin and downstream target CCND1 was not significantly affected by sost-deficiency. Further, the cell nuclear fraction of unphosphorylated (active) β-catenin was less in the *sost* KO IVD than in WT IVD. Lastly, Wnt signaling inhibitors dkk1, a downstream target of Wnt signaling,^18^ along with Wnt ligand inhibitor, *sfrp4*, was highly upregulated in *sost* KO IVD. Compensation of Wnt signaling occurs in the IVD^19^ and other tissues.^11,22^ However, it is unclear if this compensation would occur in degenerated IVD or osteoporotic vertebrae.

### Cell Phenotype

Cells of the NP, or notochordal cells, can be marked by *krt19*, *foxa2*^28^, and *gdf5*^34^, among several other genes^43^ and have a chondrocytic phenotype, expressing collagen 2 and aggrecan.^30^ Here, sclab injection decreased *col2* gene and protein expression but maintained *aggrecan* and *Krt19* expression preserving the cell phenotype. While the KO decreased col2 gene and protein expression as well, notochordal expression was reduced (decrease in *Foxa2* and *Gdf5*) as well as aggrecan. A compensatory inhibition of Wnt signaling in response to *sost* deficiency may have induced the replacement of notochordal cells by CLCs as determined by gene and protein expression of Osterix/Sp7.^20^ Injection of scl-Ab decreased osterix gene and protein expression while the KO increased the gene and protein expression of markers of CLCs (Osterix and BGLAP (Osteocalcin)). Osterix is a marker of pre-hypertrophic chondrocytes (and preosteoblasts) and the expression of osterix and BGLAP can induce and accelerate degeneration.^19,43^ The cellular shift to chondrogenesis in *sost* KO IVD is inconsistent with increased Wnt signaling,^19^ and rather, consistent of the compensation and corroborated by the moderate correlation between tail IVD degeneration and the protein cell expression of osterix. ^20^

This indicates that while these two modalities share some of the same results, a life-long deficiency of *sost* can trigger a compensatory mechanism and should be avoided in IVD with high populations of early notochordal cells, to avoid chondrogenesis resulting in dramatic changes to the IVD. Overall, these data suggest that suppression of *sost*/sclerostin may improve IVD desiccation, which is a key feature of IVD degeneration,^36^ but caution must be applied to avoid compensatory chondrogenesis.

This may be of little concern in humans as IVD degeneration begins as early as 20 years of age^4^ and anti-sclerostin antibody is administered in postmenopausal women at high risk for fracture, which commonly occurs later in age.^44^

### Difference in scl-ab and dkk, Hsp, immune, and wnt signaling

The differences in binding locations also have an implication on what class of Wnt ligands these molecules will interact with. Dkk1 will inhibit both Wnt1 and Wnt3 classes while sclerostin will inhibit only the Wnt 1 class.^36^ Due to their mechanistic differences and previous studies, sclerostin is known to be a more refined, specific inhibitor of Wnt signaling while dkk1 could function as more of a break to Wnt signaling.^36^ This could help explain the differences in heat shock protein expression.

Heat shock proteins serve to regulate the IVD in response to environmental stresses^45^ and are known to increase in response to inflammation and degeneration.^46^ *Hspa1b*, a heat shock protein found in the IVD,^47^ was differentially expressed between injections. With administration of scl-Ab, *hspa1b* was tending toward significance (p=0.06), but dkk1-Ab showed no difference from the vehicle. *Hspa1a*, another heat shock protein found in the IVD,^47^ was significantly reduced by the scl-Ab injection as well as the global *sost* KO, despite the compensation.

Activation of wnt signaling is known to promote the inflammatory response in the IVD.^44^ However, with both administration of scl-Ab and in the KO, while Wnt signaling is activated, heat shock protein gene expression is decreased. Because there was no difference in heat shock protein expression from the dkk1-Ab injection, and despite Wnt signaling being activated, we believe that heat shock protein expression in the IVD is controlled by sclerostin and not by Wnt signaling itself. Below is the heat map for Wnt signaling indicating its decrease in the *sost* KO group.

## Limitations

We would like to address some of the limitations of the study. First, we used male mice in this study but expect similar effects of deletion of *sost* to female IVD.^8,19^ Additionally, we did not have any aging data. While it is known that serum sclerostin increases with age in both male and female,^46^ the impact of estrogen-deficiency and aging on the responsiveness of the IVD to deletion of *sost* is unclear and may be impaired.^19^ Secondly, qPCR was performed on entire IVDs and the effect of *sost* KO could not be differentiated by gene expression, but key outcomes were discerned by localized expression using histology. Thirdly, silencing Wnt signaling in IVD cells induces ECM degradation,^19,48^, but also promotes anti-inflammation.^49^ By contrast, activating Wnt signaling in murine IVD leads to elevated ECM production,^50^ but also promotes pro-inflammatory signals.^49^ Based on ECM production, our data suggest that Wnt signaling was increased by anti-Wnt inhibitor injection and sost deletion (at least initially), but sost deletion also induced upregulation of immune pathways. Injection of Scl-ab did not upregulate immune-related genes, therefore, the pulsatile upregulation of Wnt signaling by Scl-Ab as occurs clinically may preclude the consequences of an inflammatory response. Despite these limitations, we show that upregulation of Wnt signaling by persistent and long-term absence of *sost* hydrates the tail IVD and promotes ECM expression in lumbar IVD, but compensation of Wnt signaling in tail IVD induces mild chondrogenesis.

## Conclusion

In addition to the well-recognized attributes to bone structure, we propose that deletion of *sost* initially increases Wnt signaling, promotes proteoglycan production, and hydrates the IVD (Fig. 5). Consequently, the osmotic pressure stimulated the cells to reduce the expression of heat shock proteins. However, compensatory upregulation of Wnt signaling inhibitors in IVD subsequently suppressed Wnt signaling, thereby triggering the loss of notochordal cells and their replacement by chondrocyte-like cells. Together, these data show that the musculoskeletal benefits of anti-sclerostin-antibody romosozumab (Evenity) may extend beyond bone to improve key features of the IVD and could potentially be used as a therapeutic for IVD degeneration.

## Abbreviations

μCT: micro-computed tomography
ACAN: Aggrecan
AF: Annulus fibrosus
BGLAP: Osteocalcin
BMD: Bone mineral density
BV/TV: Bone volume/tissue volume
CLCs: Chondrocyte-like cells
COLI: Collagen type 1
COLII: Collagen type 2
COLX: Collagen type 10
DKK1: Dickkopf WNT signaling pathway inhibitor 1
ECM: Extracellular matrix
FOXA2: Forkhead box a2
GDF5: Growth differential factor 5
IAF: Inner annulus fibrosus
IHC: Immunohistochemistry
IVD: Intervertebral disc
KO: Knockout
LRP: Low-density lipoprotein receptor-related protein
MRI: Magnetic Resonance Imaging
NP: Nucleus pulposus
qPCR: quantitative Polymerase Chain Reaction
Sclerostin: Protein product of Sost
sFRP4: Selected frizzled-related protein 4
SP7/OSX: Osterix
Tb.N: Trabecular number
Tb.Th.: Trabecular thickness
WNT: Wingless-related integration site
WT: Wild-type

---

**Supplemental Figure 1.**
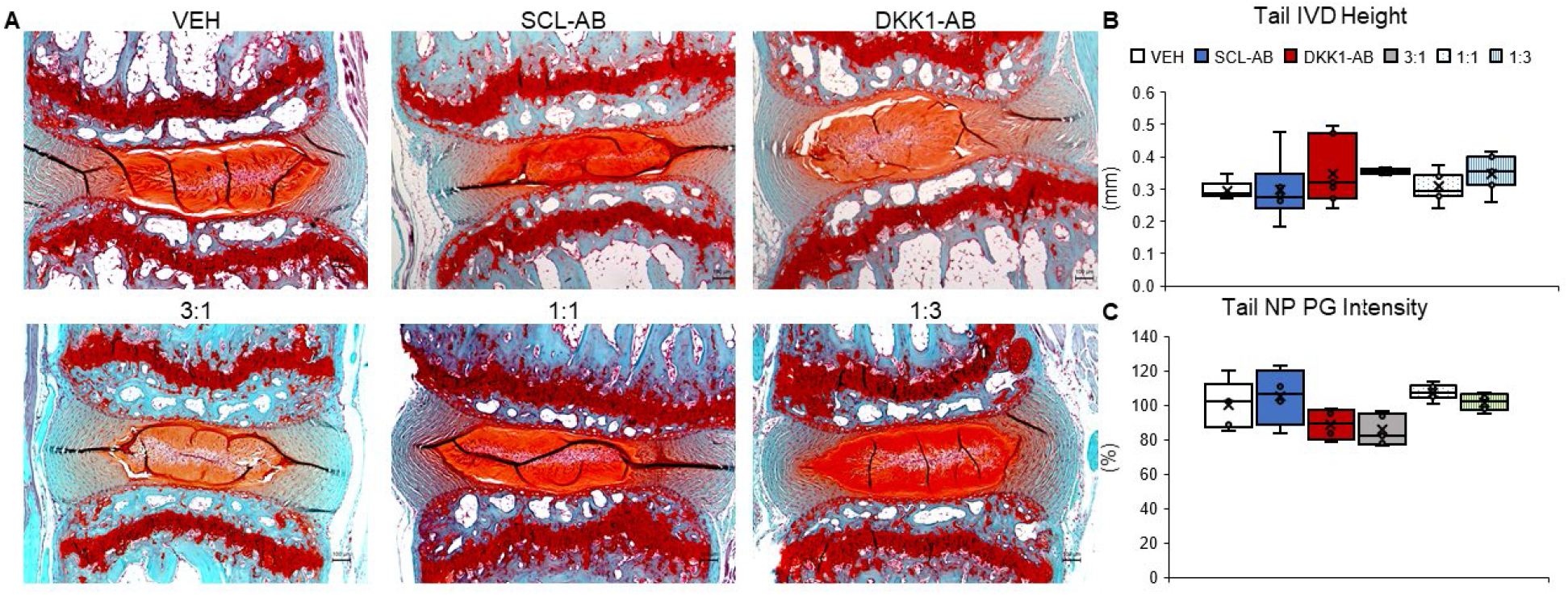
Qualitative and quantitative Tail IVD structural measurements. (A) 5X magnification images of Safranin-O and Fast Green counter stained histological sections of the tail IVD for vehicle (VEH), 25 mg/kg sclerostin-antibody injection (SCL-AB), 25 mg/kg injection of dkk1-antibody (DKK1-AB) (top row, left to right), combination injection of 18.75 mg/kg sclerostin-antibody and 6.25 mg/kg dkk1-antibody (3:1), combination of 12.5 mg/kg each of sclerostin- and dkk1-antibody (1:1), and a combination injection of 6.25 mg/kg of sclerostin-antibody and 18.75 mg/kg of dkk1-anitbody (1:3) (bottom row, left to right). (B) Quantitative measurement of tail IVD height in mm of all 6 groups. (C) Quantitative measurement of proteoglycan intensity staining in percentage of the tail IVD. Red staining indicating proteoglycan content. Scale bar is 100 μm.

**Supplemental Figure 2.**
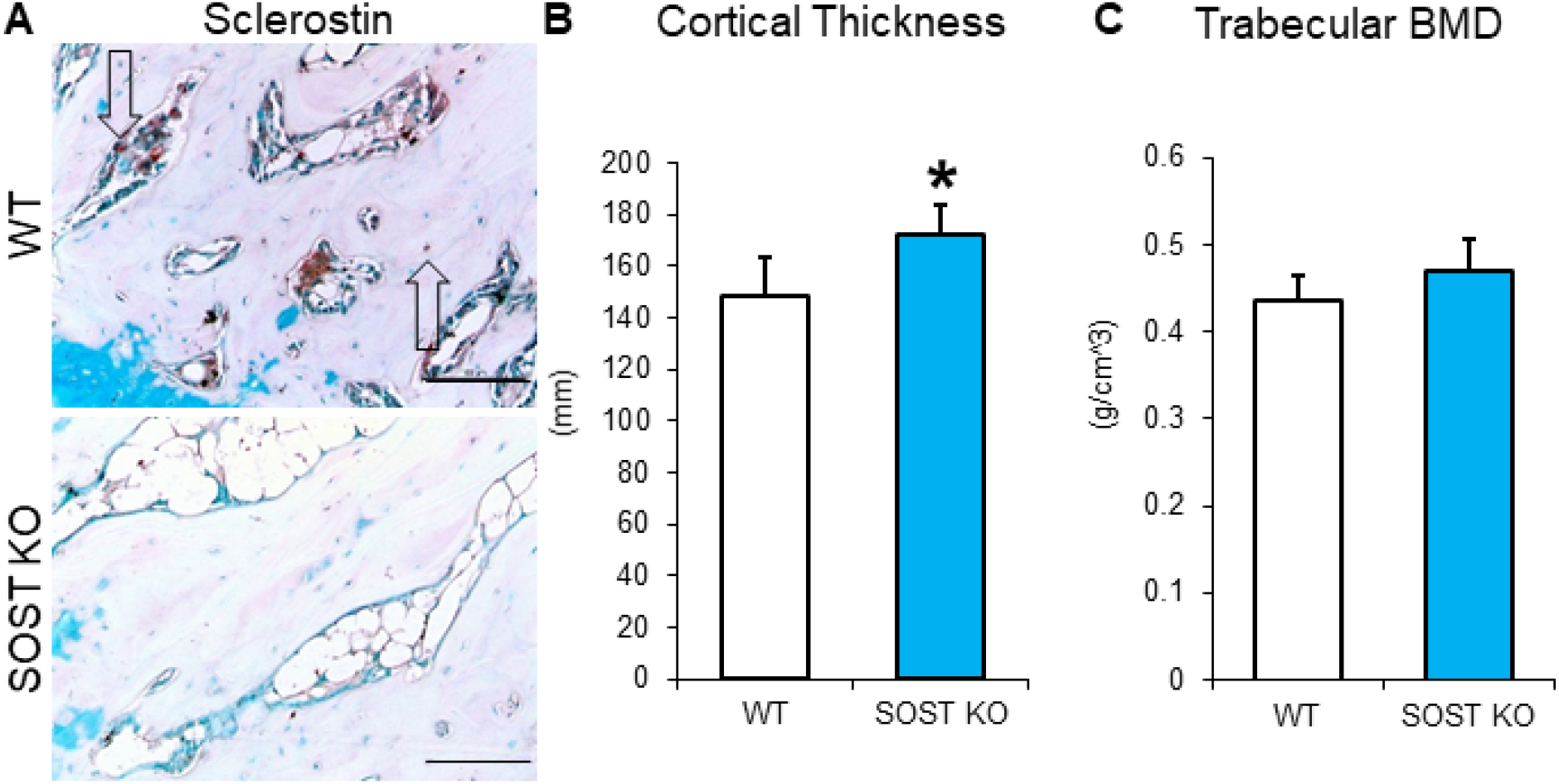
*sost* KO Bone Parameters. (A) 40X images of sclerostin stained histology slides of the vertebra. Arrows highlighting sclerostin staining in the osteocytes in the WT while there is no staining in the KO. (B) Increased vertebral cortical thickness in the *sost* KO compared to the WT. (C) Trabecular bone mineral density (BMD) for WT and *sost* KO vertebrae. Scale bar: 100um *p<0.05.

**Supplemental Figure 3.**
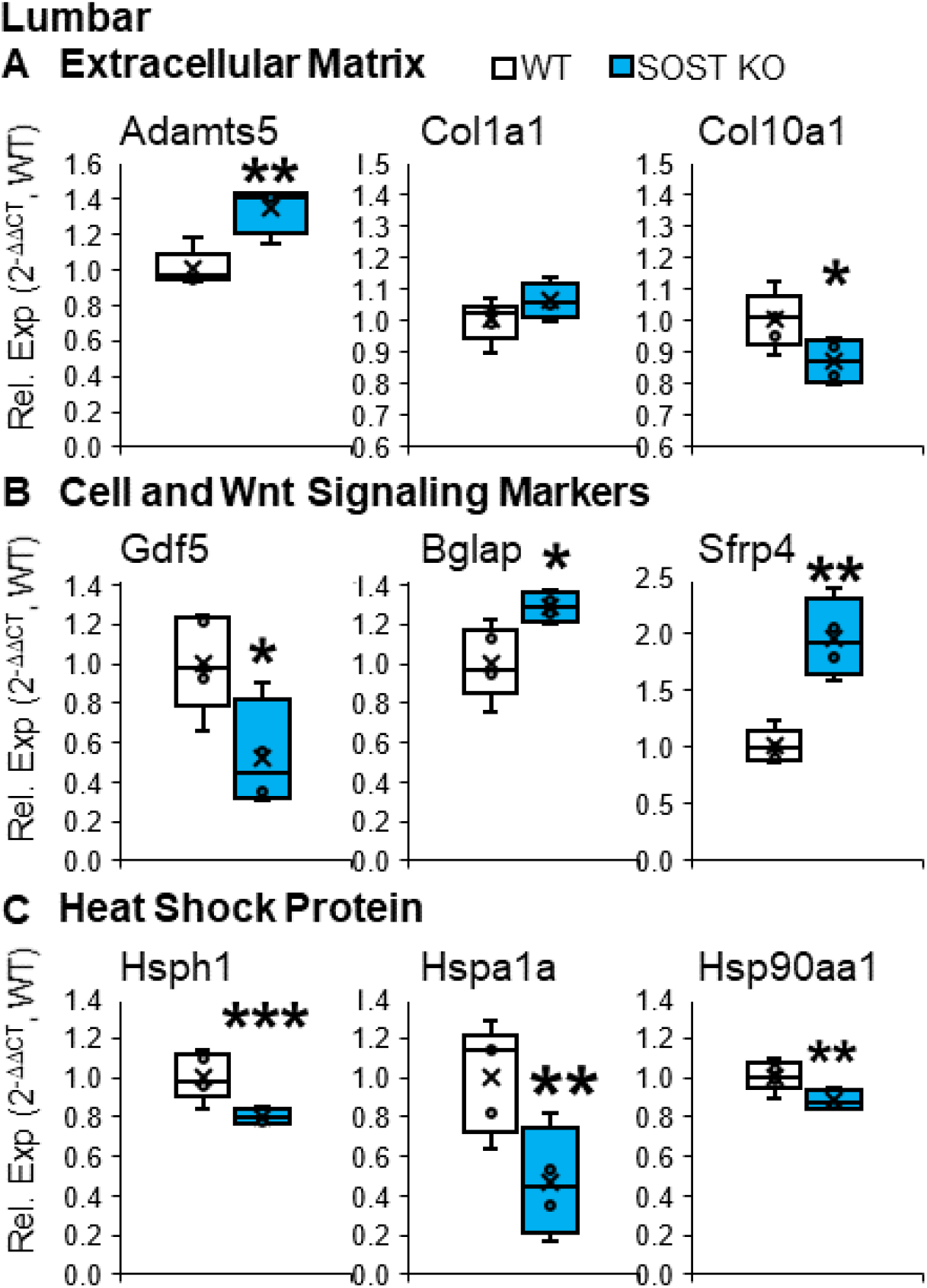
Global deletion of *sost* induced chondrogenic gene expression in the Lumbar IVD. QPCR ran on the lumbar IVD for WT and *sost* KO groups. (A) Extracellular matrix markers *adamts5* (upregulated), *col1a1* (unchanged), and *col10a1* (decreased) mirror the ecm changes in gene expression of the *sost* KO tail IVD (Fig. 7A). (B) Cell and Wnt Signaling Markers respond similarly to *sost* KO tail IVD. *Gdf5,* similar notochordal marker to *foxa2*(Fig. 7A), decreases from *sost* KO, while mature chondrocyte-like cell marker *bglap*, similar to *sfrp4* increases from *sost* KO in the lumbar just like in the tail. (C) Many heat shock proteins found in the IVD decrease from *sost* KO in the lumbar. *p<0.05, **p<0.01, ***p<0.001.

